# 𝓲-SATA: A MATLAB based toolbox to estimate Current Density generated by Transcranial Direct Current Stimulation in an Individual Brain

**DOI:** 10.1101/2020.05.28.120774

**Authors:** Rajan Kashyap, Sagarika Bhattacharjee, Ramaswamy Arumugam, Kenichi Oishi, John E. Desmond, SH Annabel Chen

## Abstract

**Background:** Transcranial Direct Current Stimulation (tDCS) is a technique where a weak current is passed through the electrodes placed on the scalp. The distribution of the electric current induced in the brain due to tDCS is provided by simulation toolbox like Realistic-volumetric-Approach-based-Simulator-for-Transcranial-electric-stimulation (ROAST). However, the procedure to estimate the total current density induced at the target and the intermediary region of the cortex is complex. The Systematic-Approach-for-tDCS-Analysis (SATA) was developed to overcome this problem. However, SATA is limited to standardized headspace only. Here we develop *individual*-SATA (𝓲-SATA) to extend it to individual head.

**Method:** T1-weighted images of 15 subjects were taken from two Magnetic Resonance Imaging (MRI) scanners of different strengths. Across the subjects, the montages were simulated in ROAST. 𝓲-SATA converts the ROAST output to Talairach space. The x, y and z coordinates of the anterior commissure (AC), posterior commissure (PC), and Mid-Sagittal (MS) points are necessary for the conversion. AC and PC are detected using the acpcdetect toolbox. We developed a method to determine the MS in the image and cross-verified its location manually using BrainSight^®^.

**Result:** Determination of points with 𝓲-SATA is fast and accurate. The 𝓲-SATA provided estimates of the current-density induced across an individual’s cortical lobes and gyri as tested on images from two different scanners.

**Conclusion:** Researchers can use 𝓲-SATA for customizing tDCS-montages. With 𝓲-SATA it is also easier to compute the inter-individual variation in current-density across the target and intermediary regions of the brain. The software is publicly available.

## Introduction

Transcranial Direct Current Stimulation (tDCS) is a noninvasive brain stimulation technique where a weak current (~ 1-2mA) is passed through anode and cathode placed on the scalp [1,2]. The placement of these electrodes (referred to as montage) governs the current flow (current density) to the target region of interest (ROI). This distribution mediates the neuro-physiological changes thereby determining the efficacy of stimulation. [3]. A number of studies have shown that cortical regions under the anode electrode will exhibit increased excitability and regions under the cathode will show decreased excitability [4]. However, there is also significant current flow in intermediary regions [5,6] with some regions having the potential to cluster the current due to tissue architecture/conductivity [7]. Research has therefore focused on computational models that could predict the pattern of current flow across the brain of an individual [5,7–12]. Realistic-volumetric-Approach-based-Simulator-for-Transcranial-electric-stimulation (ROAST) toolbox was developed to enable researchers to obtain a comprehensive overview of the current density distribution across the cortex due to a montage (Figure 1A) [13]. The output of ROAST provides the amount of current density received at every x, y, and z coordinates in the native space. Though useful, such distribution restricts an objective measurement of the amount of total current density induced at target ROI and intermediary regions in the cortex. Such objective evaluation of current peaks is necessary to determine whether a particular montage yields the expected behavioural outcome. Recently, a modified version of ROAST-target was developed, where a computer algorithm can determine the placement of electrodes such that a target coordinate is stimulated with either maximum intensity or focality [14]. This approach is mainly pertinent for “high-definition” and multi electrode configuration [14]. Pragmatically, such optimization is appropriate for those behaviors where the target region (like primary motor cortex M1) is well studied and fluctuations in behavioral outcome could be monitored through neurophysiological measures (e.g., motor/sensory evoked potential). In case of higher cognitive functions (like reading, learning, memory or speech), researchers generally tend to stimulate broader regions (e.g., Supra-marginal gyrus, Inferior frontal lobe, Prefrontal cortex), where the relationship between target region and the behavior is not clear. Thus, experimenters still depend on a subjective evaluation of the outputs from conventional modelling software. An objective quantification of total current density received at the target ROI and intermediary regions could be helpful. This is essential to reduce inconsistencies in montage selection as well [15].Systematic-Approach-for-tDCS Analysis (SATA) was developed by our group as a post-processing toolbox to estimate the current density induced in each lobe and gyri in the cortex (Figure 2) after a montage is simulated in ROAST [16]. However, SATA was limited to the standard (MNI152) built-in head model incorporated in ROAST. Here we propose *Individual*-Systematic-Approach-for-tDCS Analysis (𝓲-SATA) that could provide the current density distribution across each lobe/region after a montage has been simulated in ROAST that is applied to an individual’s brain image (i.e., T1-weighted image taken from magnetic resonance imaging).

**Figure 1.**
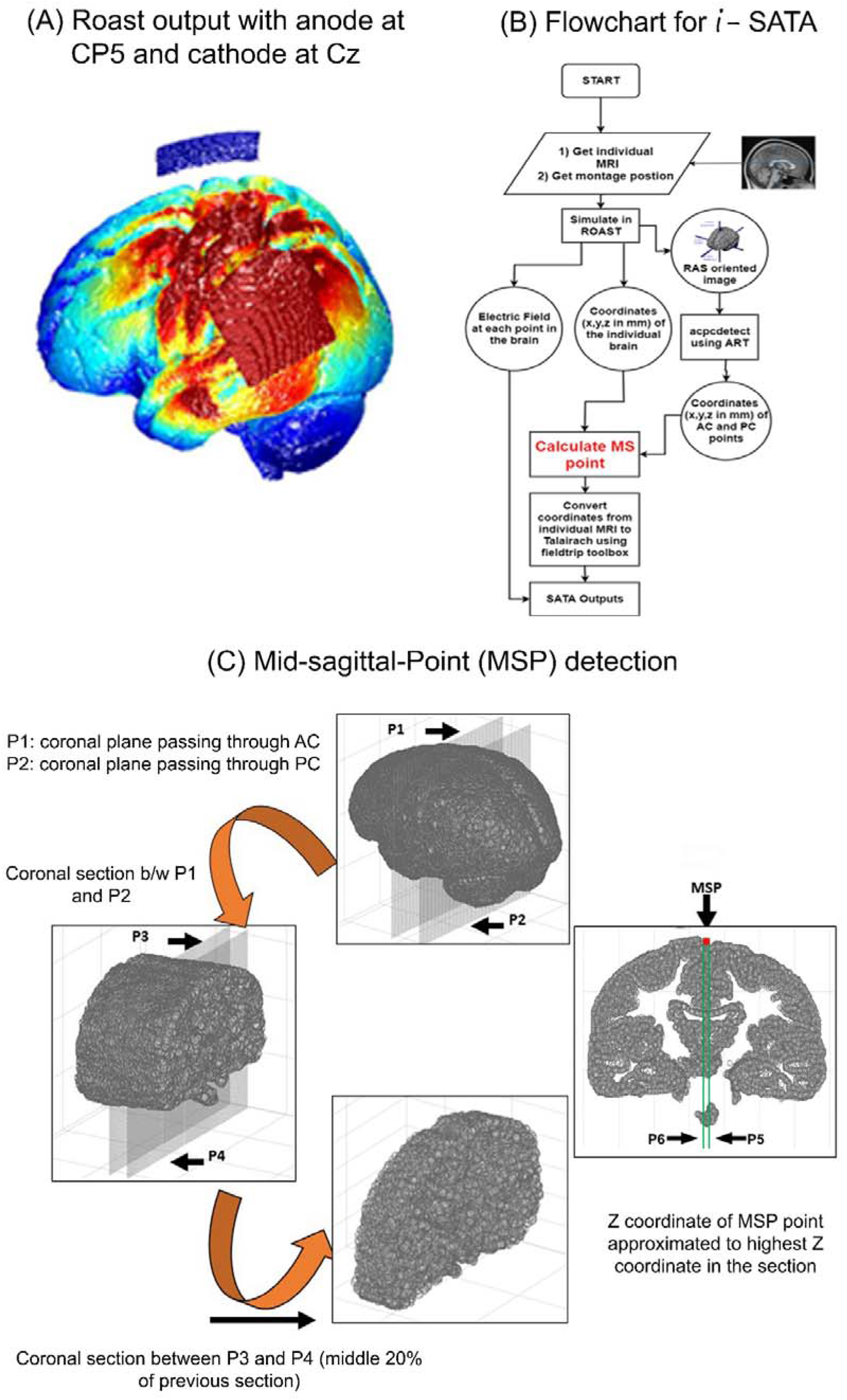
Illustration of the problem, flowchart, and the key strategy adopted for 𝓲-SATA. (A) The current density distribution over the cortex for a montage with anode at CP5 and cathode at Cz simulated in ROAST on the standard MNI-152 brain image. The amount of stimulation received by the target and intermediary areas cannot be determined objectively. (B) Flowchart displays the steps involved in 𝓲-SATA. In short, montages need to be simulated in ROAST on an individual’s T1-weighted image and 𝓲-SATA will convert the individual image to the Talairach space. For this, determining the Mid-Sagittal (MS) point of an image was important. (C) The procedure involved in determining the MS point from a brain image is explained in steps (1 to 4).

**Figure 2.**
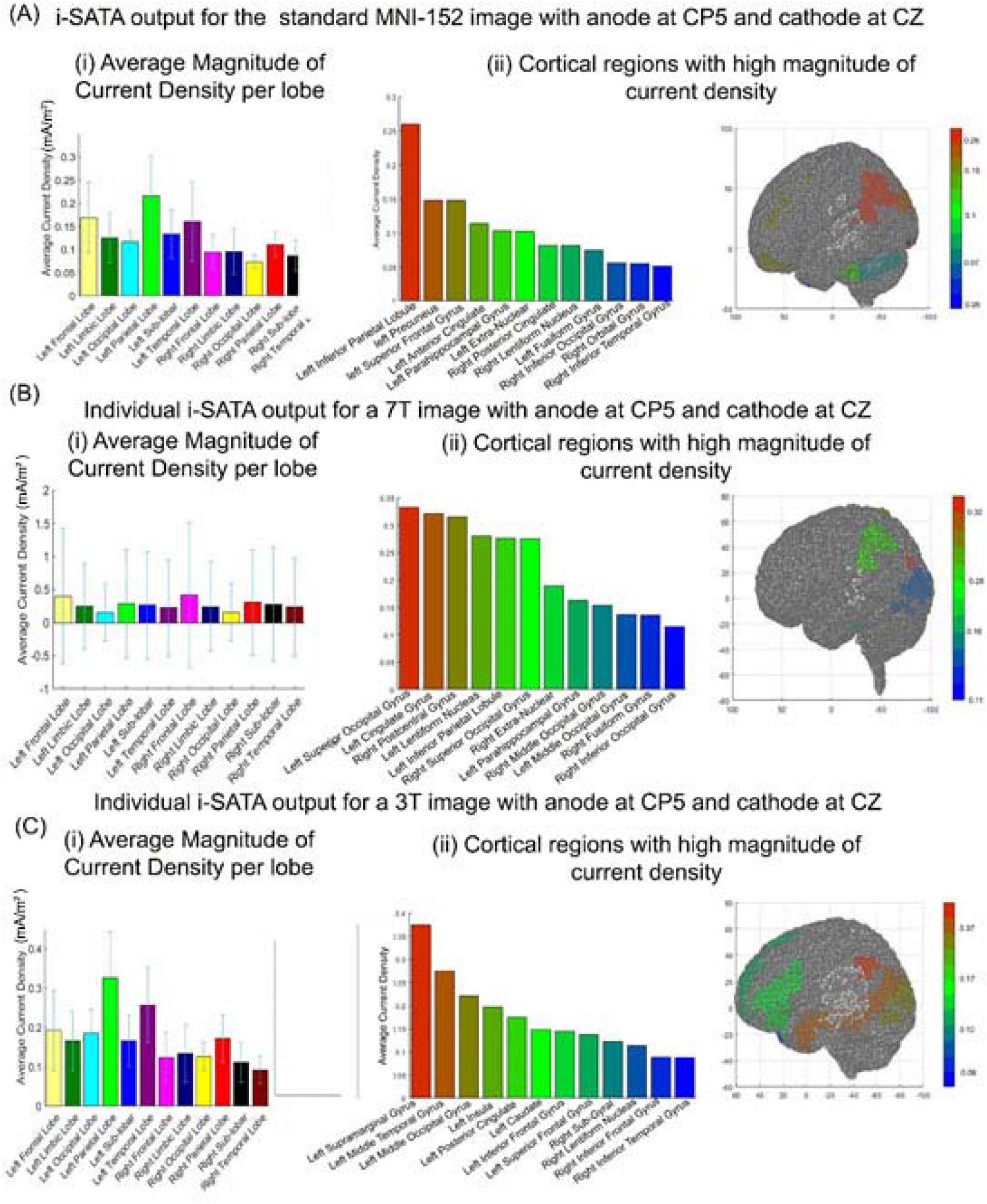
(A) shows the SATA output (MCD, CH-MCD and its distribution across the cortex) for a standard MNI-152 image for a montage with anode at CP5 and cathode at CZ with electrode size 5 × 5 cm^2^. (B) and (C) shows the SATA outputs for the same montage applied on an individual image from Data-7T and Data-3T respectively.

SATA provided two important outputs to facilitate users in choosing an appropriate montage (see methods). To obtain the outputs, the coordinates of the standard head model were mapped to Talairach space and the coordinates were allocated a cortical label. In short, coordinates belonging to a designated lobe (e.g., inferior parietal lobe, occipital lobe, etc.) were identified by the Talairach-client [17] included in SATA. Three points from the structural image namely the anterior commissure (AC), posterior commissure (PC), and mid-sagittal (MS) are necessary for conversion. AC and PC points can be identified using the acpcdetect toolbox [18], but MS was detected manually. Here we developed an algorithm that could automatically detect the MS point from the structural image of the individual brain (Figure 1B).

Individualization of SATA is important because research has shown that response to tDCS varies across individuals despite being stimulated by the same montage [19–21]. Several factors (e.g. neuronal state, anatomy, age, current intensity, etc.) lead to inter-individual variations in the induced electric field or current density [19–27]. Many computational pipelines facilitating the individualization of tDCS dosage have been developed [17, 24–27]. The main focus of these studies was to either (1) develop a realistic head model for their study [32], or (2) determine the individualized dose for specific regions of interest like M1 [33] that has specific interest amongst researchers [34]. Although the approach provided detailed information about M1 but manifestation across other ROIs were limited. With 𝓲-SATA, we foresee that the current density can be estimated at ease across the whole brain of an individual including the target and intermediary regions for any montage and thus, the estimation of inter-individual variation in the current density can also be effortlessly done.

## Methods

### Data

Two publicly available datasets were considered in the study to validate 𝓲-SATA across different MRI scanners. The first dataset with structural images (T1-weighted) of 10 healthy participants obtained from a 7-Tesla MRI scanner was taken from the Study Forrest Project (www.studyforrest.org) [35]. The dataset was referred to as ‘Data-7T’. Similarly, a second dataset (‘Data-3T’) comprising of T1-weighted images (MP2RAGE) of 5 participants imaged at a 3-Tesla MRI scanner was obtained from the Max Planck Institut Leipzig Mind-Brain-Body Dataset – LEMON (http://fcon_1000.projects.nitrc.org/indi/retro/MPI_LEMON.html) [36].

### Preprocessing with ROAST

We simulated a montage with anode at CP5 and cathode at Cz with the electrode size 5 × 5 cm^2^ and total current intensity of 2mA for all the head models. Default values of conductivity values of the tissues (white matter (default 0.126 S/m); grey matter (default 0.276 S/m); cerebrospinal fluid (default 1.65 S/m); bone (default 0.01 S/m); skin (default 0.465 S/m); air (default 2.5e-14 S/m); gel (default 0.3 S/m); electrode (default 5.9e7 S/m)) were used for each image when simulated in ROAST. Outputs of ROAST included the locations (x, y, and z coordinates) of the brain regions and the current density (mA/m^2^) value at each location. ROAST also provided the right anterior superior (RAS) oriented image of the brain. It is important to mention that such pre-processing can also be done using other freely available software. Three main inputs are needed to post-process the simulation outputs in 𝓲-SATA: 1) RAS oriented image to locate the anatomical landmarks 2) The coordinate matrix and 3) the corresponding current-density matrix that is obtained after simulation.

### Determination of anatomical landmarks

Three main anatomical landmarks are required which are AC, PC and MS point. The coordinates of AC and PC points were then obtained from the RAS-oriented brain images using acpcdetect [18]. These points were further used to determine the coordinates of the MS point.

### Determination of MS point

If we draw a line connecting the coordinates of AC and PC point, then the coordinates of MS point in the cortex will lie on the plane drawn vertical from the middle point of the line (Figure 1C). Therefore, the x coordinate of the MS point (*MS_x_*) lying at the midpoint of the line connecting the x-axis of AC point (*AC_x_*) and PC point (*PC_x_*) can be calculated as

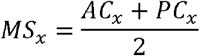

Similarly, the y coordinate of the MS point (*MS_y_*) lying at the midpoint of the line connecting the y-axis of AC point (*AC_y_*) and PC point (*PC_y_*) is

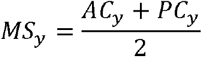

However, to determine the z coordinate of the MS point (*MS_z_*) we need to obtain the coronal section of the brain lying between the planes passing through the AC and PC points, respectively (see Figure 1C-p1 and p2). These planes serve as boundaries for the detection of the z coordinate of the MS point. We eventually narrow down the boundaries of the coronal section by drawing two more planes p3 and p4 (p3-p4) between the planes p1 and p2 such that only 20% of the total points remain in the coronal section. The threshold of 20% was based on our experience across multiple brain images and is done to reduce the computation time. Similarly, planes p5 and p6 (p5-p6) are drawn on the sagittal section surrounding the interhemispheric fissure. These two planes were at the distance of 1mm from the *MS_x_* point. This distance was chosen to confine the z-axis points to the MS plane [37]. Thus, the z coordinate of the MS point is the maximum value amongst all the points lying on the cortex in the p3-p4 coronal and p5-p6 sagittal sections. Hence, *MS_z_* is

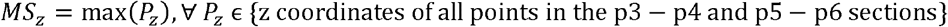

The coordinates of MS point obtained by the method were cross-verified against the MS points obtained by BrainSight^®^ (https://www.rogue-research.com/tms/brainsight-tms/). Here it is important to mention that since there is no automated method to detect MS point (based on our knowledge), the MS points from BrainSight^®^ were detected manually. The 3D Euclidean distance 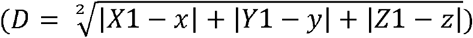 between the points manually detected by BrainSight and automatically determined by 𝓲-SATA was taken as a measure of discrepancy (error) between the two methods [16].

### SATA-based-outputs

For each montage simulated in ROAST, 𝓲-SATA extracts the location (x, y, and z coordinates) of all the points in the cortex and maps them to Talairach space. The conversion to Talairach head-space is done by inputting the detected AC, PC and MS points to the function *ft_headcoordinates* of the fieldtrip toolbox (http://www.fieldtriptoolbox.org/faq/how_are_the_different_head_and_mri_coordinate_systems_defined/) [38] and obtaining the 4×4 transformation matrix that comprises of the conversion parameters (i.e., rotational, translational, shift, and perspective, for details http://air.bmap.ucla.edu/AIR5/homogenous.html). Labeling of the coordinates to specific-cortical lobes and gyri is done by using the Talairach-client (http://www.talairach.org/) [17,39]. The magnitude of current density corresponding to each location is then used to obtain the: (1) Average magnitude of current density (MCD) received by each cortical lobe of the brain, and (2) Coordinates with high MCD (CH-MCD) in each cortical area within the lobe. It is important to mention that MCD and CH-MCD provides an estimate of the current density induced in the target and intermediary region. An ideal selection of montage for an individual should have higher values (for MCD and CH-MCD) in the targeted area, and lower values in the intermediary regions.

### Inter-individual variation

We show the inter-individual variation in the mean of the MCD and CH-MCD across the subjects in the two datasets for the simulated montage with anode at CP5 and cathode at Cz. For CH-MCD, we only highlight the important gyri in the ROI.

### Code availability

𝓲-SATA is a Linux-based-MATLAB toolbox integrating acpcdetect v2.0, fieldtrip, and Talairach Client version 2.4.3. It is operated from the command line. The toolbox can be downloaded at (LINK_TO_BE_ADDED). A reference manual is also provided for stepwise clarification.

## Results

### Cross-verification of MS point

The coordinates of the MS Points detected by BrainSight^®^ and 𝓲-SATA and the difference between the points determined by two methods are shown in Table1 for the subjects from Data-7T and Data-3T. MS points determined are in close proximity to the points obtained by BrainSight^®^. Differences between the points obtained by the two methods were calculated. The difference in-x-direction was (1.58 ± 0.85), y-direction was (1.74 ± 1.13), and in z-direction was (2.39 ± 0.88) respectively. The overall discrepancy between BrainSight^®^ and 𝓲-SATA was (1.76 ± 0.49). A fairly good degree of reliability was found between BrainSight^®^ and 𝓲-SATA measurements. The average intraclass correlation coefficient (ICC) was 0.71 with a 95% confidence interval from 0.44 to 0.86 (F(29,58) = 3.38, p < 0.0001). We cross-validated this using a fieldtrip function where MSP points were also manually located. Results were concordant with ICC = 0.77 with a 95% confidence interval from 0.52 to 0.93 (F(29,58) = 3.65, p < 0.0001).

**Table 1.** shows the location (x, y and z coordinates) in mm for the MS point determined using 𝓲-SATA and BrainSight^®^. Differences (absolute) in the points detected by the two methods is also highlighted.

| Subject | MS Point manually detected using Brainsight (in mm) |  |  | MS Point auto-detected in *i*-SATA (in mm) |  |  | Difference between BrainSight and *i*-SATA (absolute in mm)) |  |  | Discrepancy between BrainSight® and *i*-SATA |
| --- | --- | --- | --- | --- | --- | --- | --- | --- | --- | --- |
|  | X1 | Y1 | Z1 | X | Y | Z | dx | dy | dz | error |
| 1 | 104 | 138.34 | 245.27 | 104.65 | 138.5 | 245.51 | 0.65 | 0.16 | 0.24 | 0.71 |
| 2 | 105 | 133.83 | 230.51 | 106.1 | 133.05 | 229.66 | 1.1 | 0.78 | 0.85 | 1.59 |
| 3 | 110 | 141.17 | 233.98 | 109.8 | 139.5 | 235.5 | 0.2 | 1.67 | 1.52 | 2.26 |
| 4 | 105 | 140.72 | 247.83 | 104.6 | 140.4 | 246.52 | 0.4 | 0.32 | 1.31 | 1.40 |
| 5 | 105 | 140.82 | 241.28 | 103.7 | 140.5 | 242.52 | 1.3 | 0.32 | 1.24 | 1.82 |
| 6 | 107 | 140.82 | 241.58 | 107.75 | 140.45 | 242.57 | 0.75 | 0.37 | 0.99 | 1.29 |
| 7 | 112 | 135.36 | 225.57 | 110.65 | 136.55 | 226.54 | 1.35 | 1.19 | 0.97 | 2.04 |
| 8 | 102 | 127.05 | 230.86 | 103.05 | 125.95 | 232.55 | 1.05 | 1.1 | 1.69 | 2.27 |
| 9 | 103 | 130.63 | 233.61 | 101.8 | 131.5 | 234.55 | 1.2 | 0.87 | 0.94 | 1.75 |
| 10 | 111 | 129.89 | 226.67 | 112 | 130.85 | 225.56 | 1 | 0.96 | 1.11 | 1.77 |
| 11 | 96.89 | 125.78 | 217.42 | 98.8 | 126 | 218.51 | 1.91 | 1.22 | 1.09 | 2.51 |
| 12 | 95.8 | 132.5 | 236.74 | 96.9 | 133.5 | 237.52 | 1.1 | 1 | 0.78 | 1.67 |
| 13 | 103.2 | 140.27 | 241.78 | 102.8 | 139.5 | 241.52 | 0.4 | 0.77 | 0.26 | 0.90 |
| 14 | 107.4 | 140.82 | 241.58 | 107.75 | 141.45 | 242.57 | 0.35 | 0.63 | 0.99 | 1.22 |
| 15 | 99.89 | 127.78 | 213.12 | 99.8 | 126.43 | 214.51 | 0.09 | 1.35 | 1.39 | 1.93 |

### SATA-based outputs and inter-individual variation

After the montages are simulated in ROAST, the total time (per subject) taken by 𝓲-SATA on a computer (with Ubuntu 18.04 operating system and 8GB RAM) to compute the MCD, and CH-MCD is ~ 5 min each. Figures 2 shows the MCD, CH-MCD and the distribution across the brain regions for the montage with anode at CP5 and cathode at CZ with electrode size 5 × 5 cm^2^. Figure 2A shows the 𝓲-SATA output for standard MNI-152 brain image. Figure 2B and C show the 𝓲-SATA outputs for an individual from Data-7T and Data-3T. The mean and standard deviation of the current density values across the regions of interest can be visualized. Another example for simulation output for montage with anode at TP7 and cathode at supraorbital region is shown in Fig 1S in the supplementary.

Figure 3A shows the inter-individual variation in the average value of MCD and CH-MCD for Data-7T whereas 3B shows the same for Data-3T. Only two regions of interest (Inferior Parietal Lobule and Middle-Inferior Temporal Gyrus) were shown for inter-individual variation in CH-MCD in Fig 3A-ii and 3B-ii. However, a user can select many more areas as well. The green dots in Fig 3A and B represent the corresponding values when the montages were applied to the standard (MNI-152) head model in ROAST and postprocessed in 𝓲-SATA. We show the values of standard head model as we think it will help readers to understand why inter-individual variation could an important issue and the need to use 𝓲-SATA to estimate current density across the cortex of each individual. There can be subjects who have high values of MCD and CH-MCD in the intermediary regions and low in the target region.

**Figure 3.**
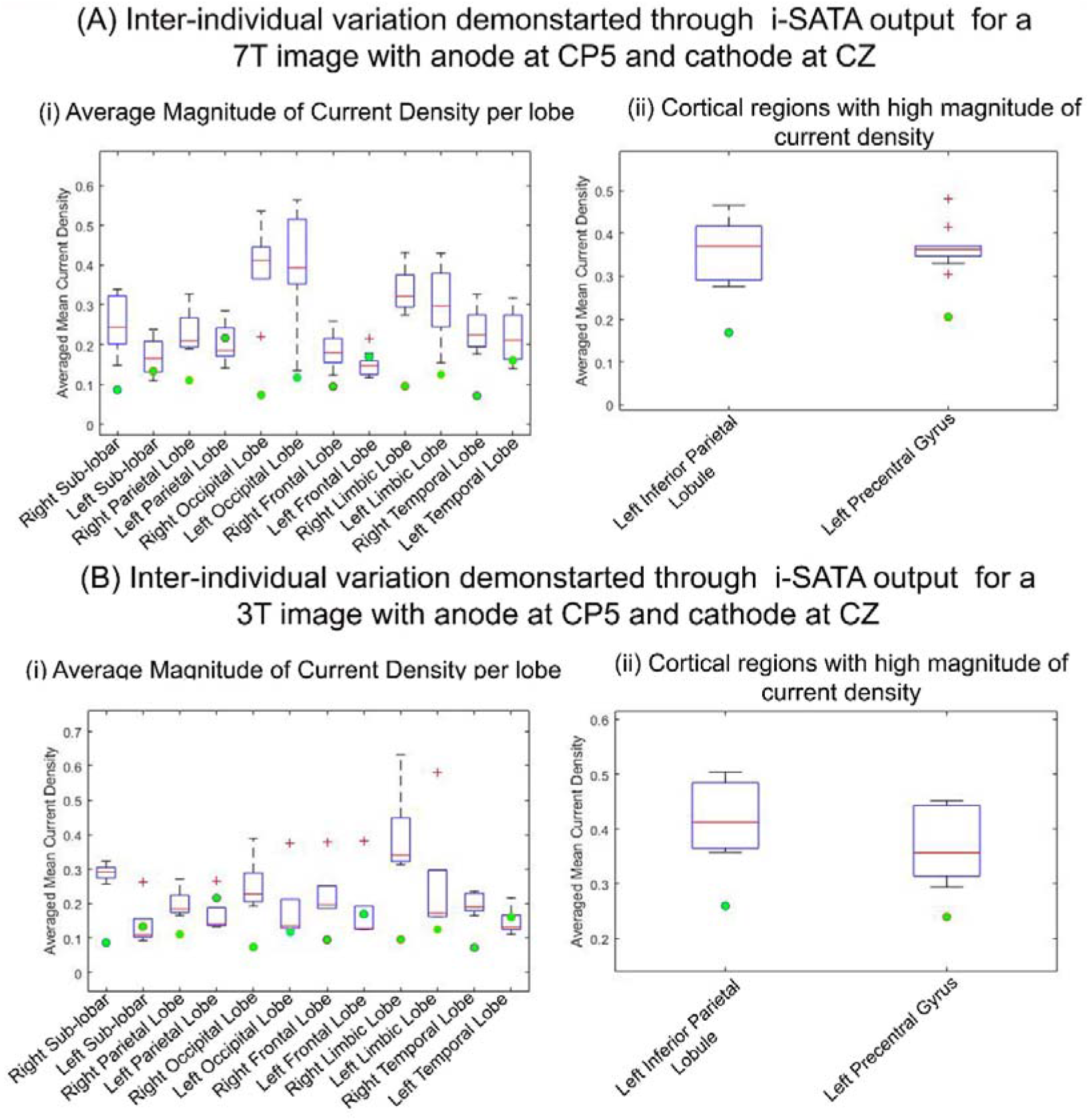
(D) and (E) shows the inter-individual variation of SATA based outputs (MCD and CH-MCD) for Data-7T and Data-3T, respectively. The green dots represent the corresponding values for the montage simulated on the standard (MNI-152) head model.

The maximum current density is found to be formed in the Left Parietal lobe for the standard head model (MNI-152) as well as for individual images taken from Data-7T and Data-3T respectively. Similarly, the target ROI (inferior parietal lobule) is found to receive substantial stimulation across the standard and individual head models. However, the variations across the intermediary regions such as the left occipital gyrus and the left middle temporal lobe are noteworthy. The inter-individual variation in the MCD and CH-MCD shows that the target lobe and gyri receive a similar amount of stimulation as received by the standard head model (green dots in the range of the error box), however, the variation in other intermediary regions like occipital, frontal and limbic lobes are evident (green dots approximately outside the range of the error box).

## Discussion

In this paper, we developed the toolbox 𝓲-SATA that provides the estimate of current density induced at each cortical lobes and intermediary regions of an individual brain administered with tDCS. To this aim, we also developed a way to detect the MS point from AC and PC points. MS points computed were in close proximity to the points traced manually using BrainSight^®^. We tested 𝓲-SATA on T1-weighted images obtained from scanners of different strengths. 𝓲-SATA provided estimates for MCD and CH-MCD of individuals. These estimates could aid researchers to choose stimulation parameters (like electrode placement, dosage, etc.) appropriate for an individual. Previous studies have found that montages tailored for each individual (individualized montage) could increase the electric field intensities at the target ROI [40]. Such individualized montages applied on stroke patients were found to improve the behavioural performance as well [40]. However, with such an approach the current intensity at target ROIs could not be maintained constant across all individuals. On a similar note, but from a different perspective, Evans et al., [28] put forward the view of individualized dosage wherein the current input is tuned to a selected montage. Although this procedure could maintain a fixed stimulation intensity at the target ROI, the spread of current across intermediary regions could not be controlled. Thus, irrespective of the individualized approaches defined by previous studies, researchers are lacking a toolbox that provides the information about the current intensity both at the target ROI and at intermediary regions for each individual. With 𝓲-SATA, outputs of MCD at target region and CH-MCD in the intermediary regions can be simulated. With this information, researchers will be able to make an informed decision regarding the appropriateness of stimulation parameters.

Having said that, we would like to highlight that the work of Evans et al.,[28] and Caulfield et al., [33] have added another parameter that the user can calculate with 𝓲-SATA i.e., the individualized dosage of current. We will explain this with an example for an individual head model. Suppose, the calculated MCD for the montage (CP5-CZ) is 0.25 mA/m^2^ at the target ROI (i.e. left inferior parietal lobe) when 2 mA of current is applied on the scalp. To achieve an intensity of 0.5 mA/m^2^ at the target ROI, the required dosage can be calculated as 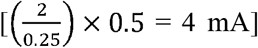 (for details, refer to [33]). We foresee this parameter strengthening the usability of 𝓲-SATA at the individual level.

With 𝓲-SATA it is easy to evaluate the inter-individual variation in MCD and CH-MCD. A user simply needs to simulate a montage in ROAST and obtain the outputs (with few MATLAB commands) in 𝓲-SATA. The complexity and computation time involved in obtaining the outputs is low. With 𝓲-SATA, we foresee that the inter-individual variation in the estimates (MCD and CH-MCD) across the target and intermediary regions of the brain can be correlated with several factors (for example-age, sex, brain anatomy, and electrode-size, shape, and placement, etc.) that cause variation in tDCS efficacy. Thus, empowering researchers with 𝓲-SATA toolbox can enhance our understanding of the various factors that influence inter-individual differences in tDCS. With that said, we would like to mention that 𝓲-SATA is not designed for pathological heads, but there are plans to add this capability in future versions.

## Conclusions

We develop the toolbox 𝓲-SATA (as an extension of SATA) that will provide estimates of current density across each cortical lobe of an individual brain. With 𝓲-SATA, it easier for users to account for the factors responsible for causing the inter-individual variation in tDCS.

## Acknowledgment

The work was supported by the NTU-JHU grant from Nanyang Technological University, Singapore. JD received additional support from NIH/NICHD grant U54 HD079123.

