## supplement for "𝓲-SATA: A MATLAB based toolbox to estimate Current Density generated by Transcranial Direct Current Stimulation in an Individual Brain"

CRADLE, SSS, LKCMedicine

Nanyang Technological University, Singapore

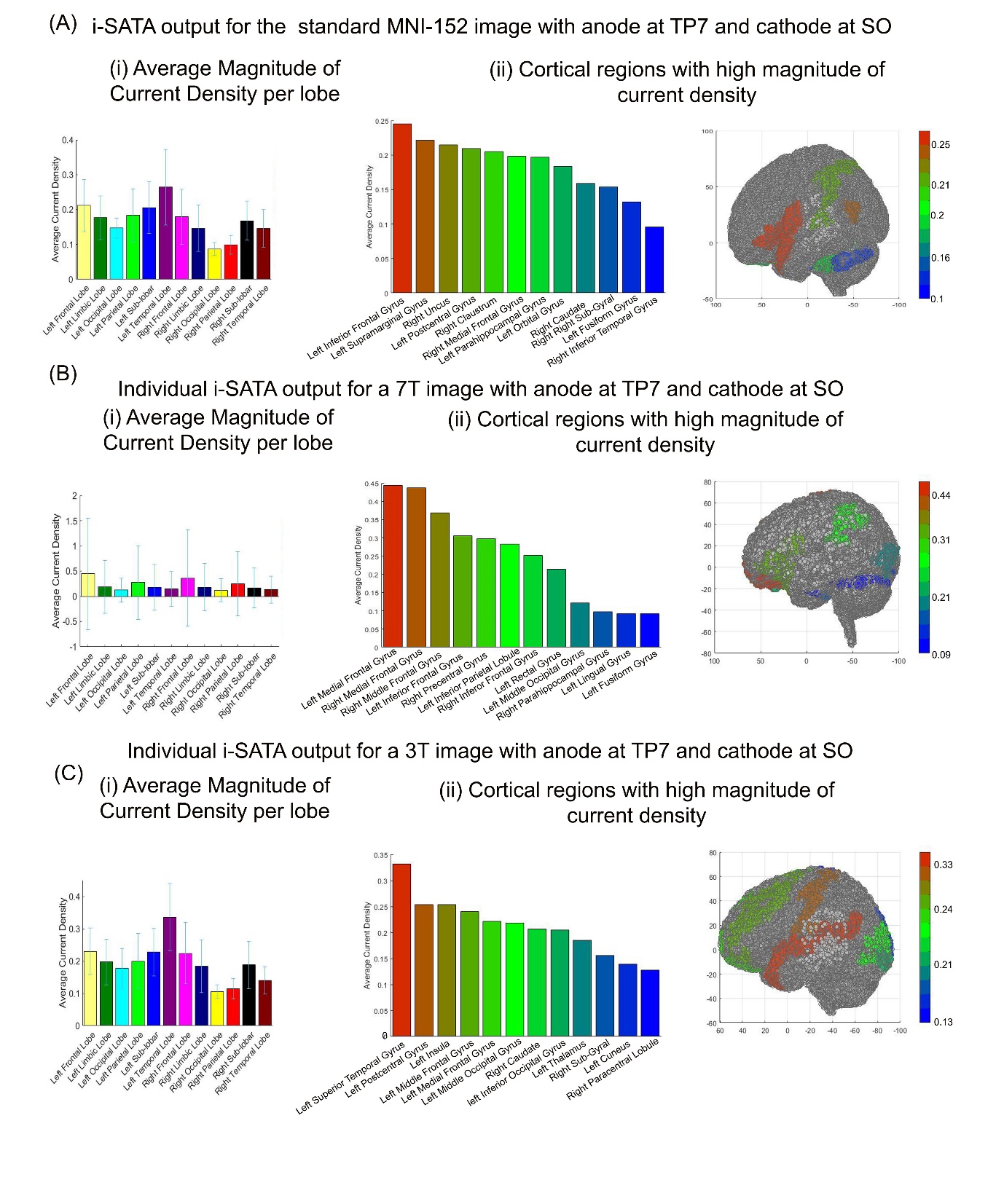

Figure 1S (A) shows the SATA output (MCD, CH-MCD and its distribution across the cortex) for a standard MNI-152 image for a montage with anode at TP7 and cathode at supra-orbital region (SO) with electrode size 5 × 5 cm^2^. (B) and (C) shows the SATA outputs for the same montage applied on an individual image from Data-7T and Data-3T respectively.

***Alternate Ways of Brain Mapping***

Here we used AC, PC and MSP points to map an individual brain space to Talairach Space using Talairach Client toolbox. Alternatevely one may also use the SPM anatomy toolbox [1] as a template. For this purpose, transformation and deformation might be attempted using SPM [2] and DARTEL [3], respectively.

***Reference***

[1]	Eickhoff S., Rottschy C., Caspers S. (2013) Tool for the integrated analysis of structure, function and connectivity: SPM Anatomy Toolbox. In: Schneider F., Fink GR (eds) Functional MRI in psychiatry and neurology. Springer, Berlin, Heidelberg

[2]	"SPM - Statistical Parametric Mapping" retreived from <https://www.fil.ion.ucl.ac.uk/spm/>

[3]	"dartel_spm documentation" retreived from <https://github.com/canlab/spm12/tree/master/toolbox/DARTEL>
